# Macrocycle screening against the C-terminal region of CHD4 uncovers its role as an interaction hub in the formation of the nucleosome remodeling and deacetylase complex

**DOI:** 10.64898/2026.06.03.728223

**Authors:** David C. Williams, Jie Ren, Torry Li, Jarrett M. Pelton, Dhanushi Dedakia, Robert K. McGinty, Gordon D. Ginder, Albert A. Bowers

**Affiliations:** Department of Pathology and Laboratory Medicine and Lineberger Comprehensive Cancer Center, School of Medicine, University of North Carolina at Chapel Hill, Chapel Hill, NC 27599; Center for Integrative Chemical Biology and Drug Discovery, Division of Chemical Biology, and Medicinal Chemistry, UNC Eshelman School of Pharmacy, University of North Carolina at Chapel Hill, Chapel Hill, North Carolina, 27599; Department of Biochemistry and Biophysics and Lineberger Comprehensive Cancer Center, School of Medicine, University of North Carolina at Chapel Hill, Chapel Hill, North Carolina, 27599; Departments of Internal Medicine, Cellular, Molecular, and Genetic Medicine, and Microbiology and Immunology, and the Massey Cancer Center, Virginia Commonwealth University, Richmond, Virginia 23298

## Abstract

The Nucleosome Remodeling and Deacetylase complex (NuRD) plays a key role in regulating hemoglobin expression in adult erythroid cells. Selectively disrupting this complex potently induces the expression of fetal hemoglobin, a proven therapeutic strategy for treating beta-hemoglobinopathies such as sickle cell anemia. In these studies, we have used mRNA display to identify small macrocyclic peptides that inhibit the interaction between two core components of NuRD, the SANT-SLIDE domain of CHD4 and the CR2 domain of GATAD2A. In addition, the screen suggested a second binding site on the CHD4 domain. Based on this observation, we hypothesized and confirmed that CDK2AP1 bound to this region of CHD4, leading us to purify and determine the structure of the ternary complex between CHD4, GATAD2A, and CDK2AP1. The results of our studies show that the SANT-SLIDE domain of CHD4 functions as a critical interaction hub in the formation of NuRD and suggest a strategy to block NuRD function for therapy.

## Introduction

Chromatin remodeling complexes reposition and evict nucleosomes, thereby modulating chromatin compaction and accessibility to play a central role in gene regulation. Disrupting the structure and function of these complexes represents a compelling therapeutic strategy for treating cancers and other diseases. In previous studies, we established that disrupting the structure of the nucleosome remodeling and deacetylase (NuRD) complex is a potential therapeutic target for restoring fetal hemoglobin expression.^1–7^ Augmenting fetal hemoglobin expression is a clinically validated and actively pursued approach to treating beta-hemoglobinopathies, including sickle cell anemia and beta thalassemia.^8–11^

Two key transcription factors, BCL11A and ZBTB7A, along with cytosine methylation, recruit NuRD to the human gamma globin proximal promoter to silence transcription and block expression of fetal hemoglobin.^12–14^ Targeted disruption of NuRD, either through genetic knockout/knockdown or inhibition of key interactions, is one of the most potent inducers of fetal hemoglobin identified to date.^1,2,5^ We have been studying the structure and function of the NuRD complex to identify targetable interactions for developing small-molecule or peptide inhibitors of complex formation.^15–18^

The canonical NuRD complex consists of 7 proteins, each of which has multiple isoforms, and can be divided into two functional and structural halves: the histone deacetylase core complex (HDAC1/2, MTA1/2/3, RBBP4/7, MBD2/3) and the large chromatin-remodeling complex (CDK2AP1/2, GATAD2A/B, CHD3/4).^19–21^ These sub-complexes are bridged by a coiled-coil interaction between the C-terminal coiled-coil domain of MBD2/3 and the conserved region 1 (CR1) coiled-coil domain of GATAD2A/B.^15^ Previous work has established that conserved region 2 (CR2) of GATAD2A/B interacts with the C-terminal region of CHD4.^16^ However, the molecular details of this critical interface remain unknown. Notably, knockdown of CHD4 strongly induces fetal hemoglobin^5^ and mutations in the C-terminal region of this protein in erythroid progenitor cells do not reduce cell fitness but potently induce fetal hemoglobin expression.^22^ The latter suggests that selectively targeting the C-terminal region of CHD4 may induce fetal hemoglobin expression to therapeutic levels without significant side effects.

In these studies, we set out to identify inhibitors of the GATAD2A-CHD4 interaction as molecular probes and potential therapeutic leads. Based on an AlphaFold 3 (AF3) model of this interaction, we performed an mRNA-display screen for macrocyclic peptide binders to the CHD4-C1bC2a SANT-SLIDE domain. Modeling the top-enriched peptide sequences bound to CHD4-C1bC2a using AF3 highlighted two distinct binding interfaces and guided the identification of competitive inhibitors of the interaction with GATAD2A. The most potent inhibitors share a common motif with an i, i+4 crosslink that stabilizes a short helix. In addition, we identified a peptide that binds CHD4-C1bC2a but does not competitively inhibit CR2 binding, consistent with binding to the second interaction site on the SANT-SLIDE domain. This result led us to explore whether CDK2AP1 bound to this region of CHD4, and ultimately to determine the structure of a stable ternary complex between CHD4-C1bC2ab, GATAD2AC-CR2, and CDK2AP1. This structure details, for the first time, the role that CDK2AP1 plays in the formation of NuRD, suggesting how it may regulate the complex’s structure and function. Hence, we have utilized a combination of machine learning, mRNA display, and structural biology to develop novel molecular probes and gained new insights into the formation of NuRD.

## Results

### GATAD2A-CR2 domain binds to CHD4-C1bC2a

In recent studies, Zhong *et al*. determined the structure of the C1b-C2a region of CHD4, revealing that it forms a SANT-SLIDE domain that binds DNA and regulates chromatin remodeling activity.^23^ An AF3 model indicates that this same region forms a complex with the GATAD2A-CR2 domain, and that this interaction involves extensive contacts with a short central helix in the CR2 domain (residues 383-401). To validate this interaction, we measured binding using a fluorescence polarization assay, which showed that the isolated helix (CR2-helix) binds to CHD4-C1bC2a with high affinity (K_D_ = 9 ± 1 nM). Likewise, an intracellular bioluminescent resonance energy transfer assay (NanoBRET) shows that the central helix and Zn finger of CR2 (individually and together) interact with CHD4-C1bC2ab in cells.

### mRNA display selection of macrocyclic peptides that bind CHD4-C1bC2a

Having established that the GATAD2A-CR2 domain binds to CHD4-C1bC2a, we sought to identify macrocyclic peptide inhibitors of this interaction by mRNA display. We deployed a library comprised of two different variable regions of either 10 or 11 amino acids, flanked by a pair of cysteine residues that could be used as handles for chemical cyclization. This library was transcribed into RNA, linked to puromycin, translated, reverse transcribed, and then cyclized with *meta*-dibromoxylene (*m*-DBX). Given the presence of a charged, DNA-binding region in CHD4-C1bC2a, we used a modified selection protocol wherein free, biotinylated CHD4-C1bC2a was incubated in solution together with library, BSA, and ss-DNA (the latter two as blocking agents to prevent the non-specific adsorption of target to the mRNA/cDNA library tag), and the resultant CHD4-C1bC2a:mRNA-display-peptide complexes were captured with immobilized, magnetic streptavidin beads. cDNA was then eluted, amplified, and resubmitted to additional iterations (see Materials and Methods). After five rounds of selection, we observed a robust recovery of the input library (**Figure 1A**), suggesting enrichment of specific binder sequences, and the cDNA was sent for Next Generation Sequencing (NGS).

**Figure 1.**
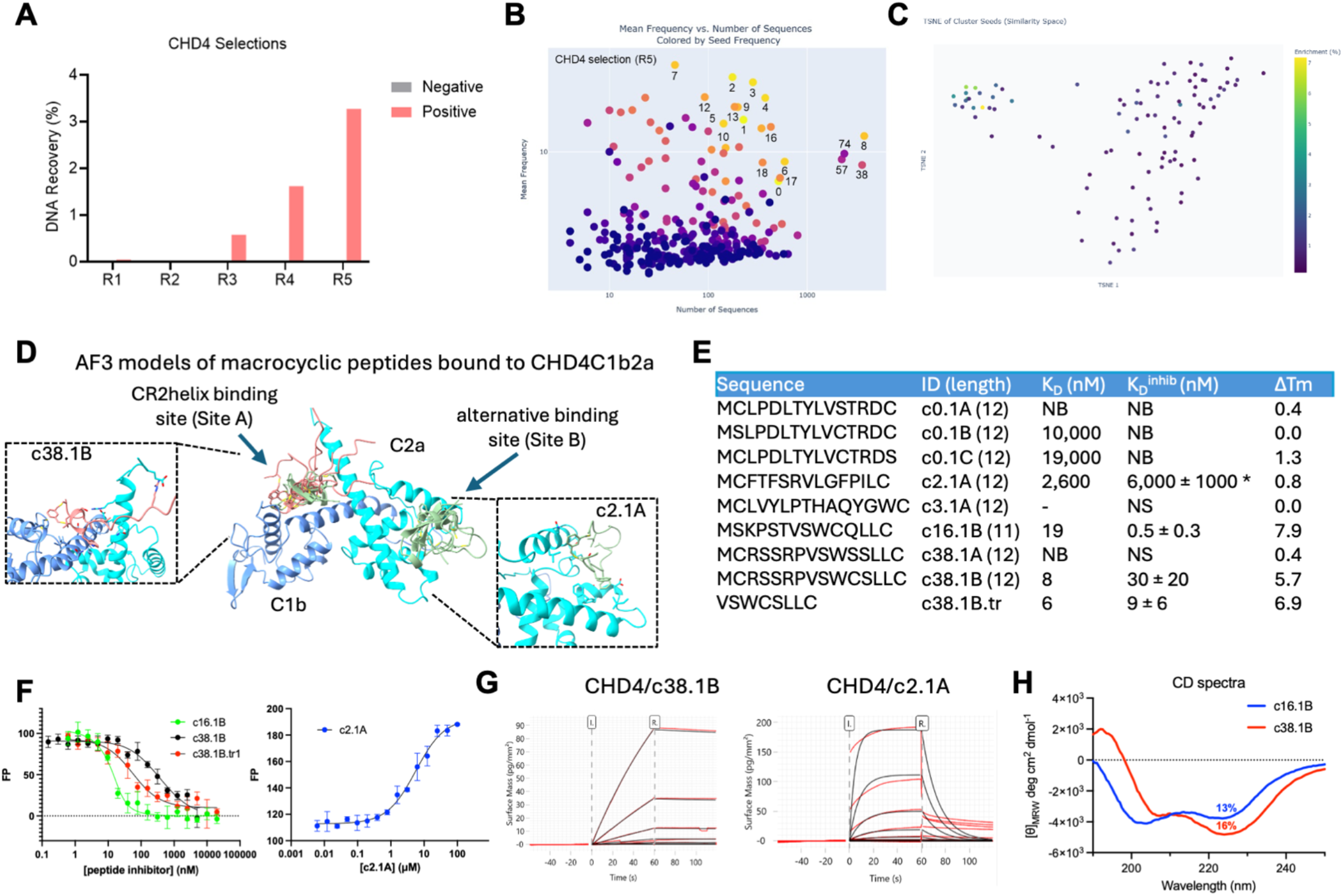
mRNA selection and characterization of peptides that bind CHD4-C1bC2a. (**A**) PCR analysis shows good recovery of the input library after five rounds of selection. Peptide sequences from round five of the selection were clustered by chemical similarity using Morgan fingerprints and plotted (**B**) based on the mean sequencing frequency of peptides within the clusters versus the number of similar peptides grouped into the clusters and (**C**) in chemical space, based on similarities between the clusters themselves, on a t-SNE plot. (**D**) AF3 modeling of top-enriched macrocyclic peptides reveals two potential binding sites. Peptides cyclized between the cysteine residues in the WCXLLC motif (light coral) are predicted to bind to Site A, while most of the remaining peptides (light green) are predicted to bind to Site B of the CHD3-C1bC2a SANT-SLIDE domain (C1b –cornflower blue, C2a – cyan). (**E**) The amino acid sequences for the top enriched peptides for each cluster are listed with the measured binding affinities as determined by GCI (K_D_) and FP competition assay (K_D_^inhib^) and the measured change in melting temperature for CHD4-C1bC2a upon binding each peptide (ΔTm) as determined by nanoDSF. (**F**) The results of fluorescence polarization competitive inhibition assay are plotted for peptides that competitively displace the CR2-helix (left panel) and the peptide (c2.1A) that non-competitively binds to CHD4-C1bC2a (right panel). (**G**) GCI binding analyses are plotted for c38.1B (competitive inhibitor) and c2.1A (non-competitive binder) binding to CHD4-C1bC2a. (**H**) Circular dichroism spectra are shown for two peptides cyclized between the cysteine residues in the WCXLLC motif (c16.1B and c38.1b) with calculated helical content.

We employed a two-step approach to analyze sequencing results. In the first step, sequences were clustered according to chemical similarity using Morgan fingerprints. Clusters were visualized in two different ways: 1) based on the mean sequencing frequency of peptides within the clusters versus the number of similar peptides grouped into the clusters (**Figure 1B**), and 2) in chemical space, based on similarities between the clusters themselves, on a t-SNE plot (**Figure 1C**). The most enriched sequences fall into different clusters that are quite distinct in chemical space (**Figure 1B**, clusters 0-7). However, there is also a large set of clusters that are comparably enriched and grouped closely in chemical space (**Figure 1B**, clusters 8, 38, 57, 74). All peptides in this latter set exhibit a short and strictly conserved C-terminal (WCXLLC) motif, opening the possibility of an alternative ring topology from cyclization of *m*-DBX onto the internal Cys residue. To further analyze this data and prioritize peptides for synthesis, we used AF3 to predict peptide binding to CHD4-C1bC2a. As shown in **Figure 1D**, the resulting models suggest two distinct binding sites for the selected peptides. Many of the topmost enriched peptides are predicted to bind to a unique site on a relatively flat surface of the C2a domain (c2.1A in **Figure 1D**), while the subset containing the (WCXLLC) motif (clusters 8, 16, 38, 57, 74) is predicted to bind to the same region as the CR2-helix (c38.1B in **Figure 1D**).

We chose to synthesize several of the peptides predicted to bind to Site A (CR2-helix binding site, representative clusters 16 and 38), as well as some predicted to bind Site B (unknown binding site, representative clusters 0, 2, and 3). For sequences containing an internal cysteine, we synthesized the three different DBX-crosslink possibilities, replacing the remaining cysteine with a serine (e.g., peptides c0 and c38 in **Figure 1E**). These compounds were first tested for their ability to inhibit CR2-helix binding to CHD4 in a competitive fluorescence polarization assay. In these experiments, the WCXLLC-containing peptides proved to be potent, low-nanomolar inhibitors, when the DBX bridge was incorporated between the two C-terminal cysteine residues (**Figures 1E, F**); other ring topologies of these peptides did not exhibit appreciable binding. We also synthesized a truncated version of this family to assess whether the WCXLLC motif alone was sufficient for binding. The resulting peptide (c38.1B.tr) likewise competitively binds to CHD4-C1bC2a with similar affinity (**Figures 1E, F**). In contrast, none of the predicted Site B binders inhibited CR2-helix binding; however, a cyclic variant of a top hit peptide from cluster 2 bound non-competitively with low micromolar dissociation constant (**Figures 1E, F**); here, polarization increases with the addition of peptide, consistent with binding to an alternative site on CHD4-C1bC2a.

We further measured binding by grating-coupled interferometry (GCI) and nano-differential scanning fluorometry (nanoDSF; Nanotemper Monolight NT.115), which confirmed that peptides bound CHD4-C1bC2a with affinities in good accord to the FP assay (**Figures 1G**). Similarly, the nanoDSF results show that changes in observed melting temperature (ΔT_m_) upon peptide binding correlate with the measured binding affinities, with the high-affinity peptide inhibitors increasing the T_m_ by up to 7.9 °C (**Figure 1E**). Notably, an i, i+4 crosslink with DBX is known to stabilize helix formation in short peptides. This observation suggests that the selected WCXLLC-motif may form a short helix stabilized by the DBX crosslink, allowing it to interact with the same cleft occupied by the CR2-helix. Indeed, we measured circular dichroism for two peptides with crosslinked WCXLLC motifs (**Figure 1H**), confirming that these peptides exhibit significant helical content in isolation. Of note, we have not identified any peptides that bind more avidly to the alternative site (Site B), suggesting that the physical characteristics of this site are not favorable for tight association.

### Second binding site in CHD4-C1bC2ab

The screen results and modeling highlight a second binding site on CHD4-C1bC2a. This observation raises the question of whether a native protein binds to this site. Of the remaining known components in NuRD, only CDK2AP1 has been shown to bind directly to CHD4. While often overlooked, CDK2AP1 was identified as a core component by the Vermeulen group.^24^ In addition, crosslinking analysis showed that CDK2AP1 bound to the C2b region of CHD4.^25^ An AF3 model indicates that the binding involves the C2b region of CHD4 (residues 1800-1912). We confirmed this direct interaction between CHD4-C1bC2ab and CDK2AP1 by NanoBRET (**Figure 2A**).

**Figure 2.**
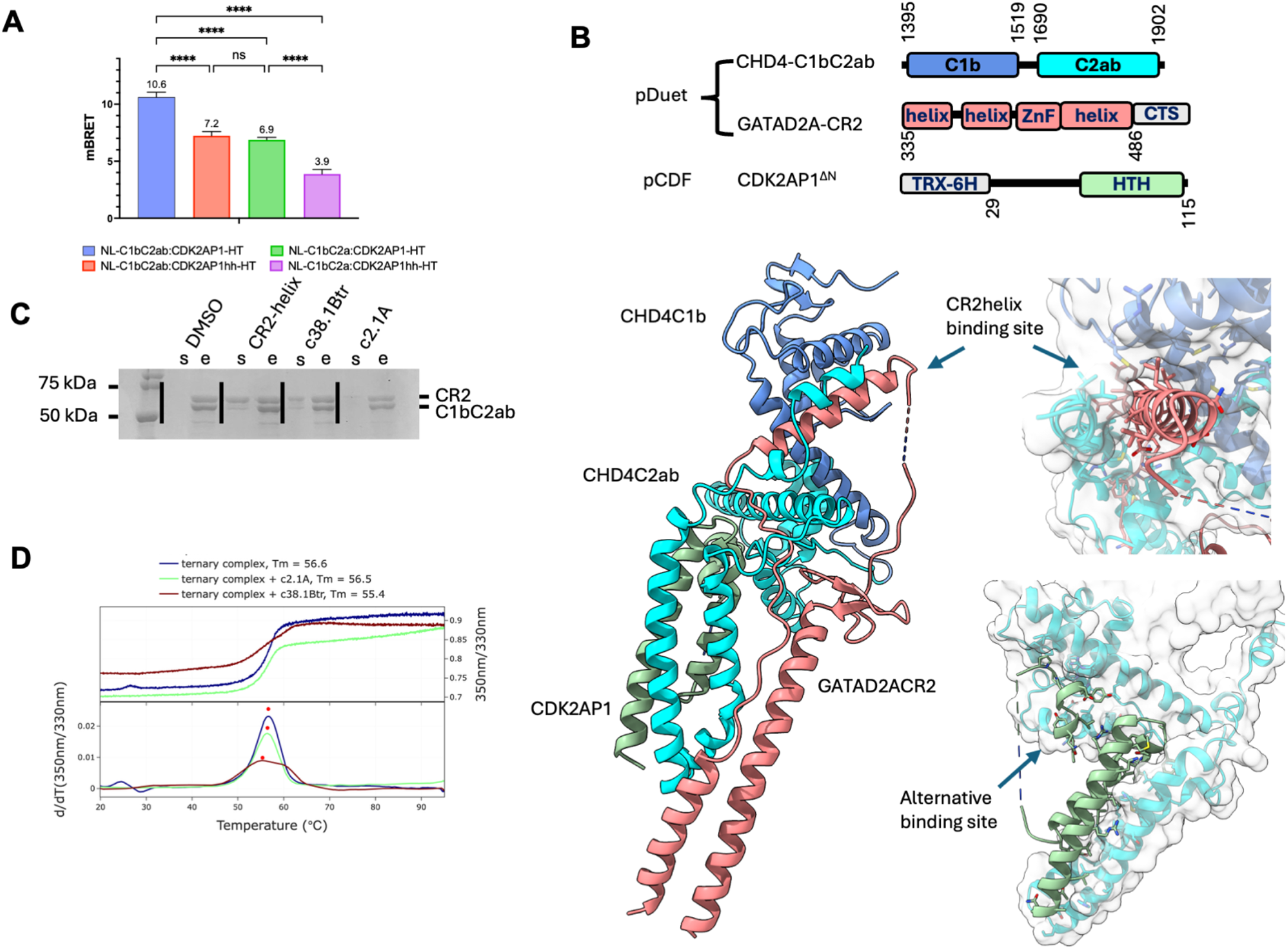
Ternary complex formation and structure. (**A**) In cell bioluminescent energy transfer analysis (NanoBRET) shows that CHD4-C1bC2ab binds to CDK2AP1. Maximum binding requires the CHD4-C2b region (C1bC2ab vs C1bC2a) and the unstructured N-terminal region of CDK2AP1 (CDK2AP1 vs CDK2AP1hh). (**B**) The crystal structure of the ternary complex was determined by co-expressing all three protein constructs in pDuet and pCDF as depicted. A ribbon diagram and transparent surface representation of the resulting structure shows how GATAD2ACR2 (light coral) forms a loop that binds at the interface between the CHD4-C1b (cornflower blue) and CHD4-C2ab (cyan) domains and interacts with the long CHD4-C2b helix hairpin. CDK2AP1 (light green) binds to the opposite face of the CHD4-C2b helix hairpin while its unstructured N-terminal region folds into a helix and binds to the alternative binding site (Site B) on CHD4-C2a. (**C**) A polyacrylamide gel of the on bead displacement assay shows that the CR2-helix and c38.1Btr peptides displace the CR2 domain from the binary complex while c2.1A does not. Lanes labeled “s” are the supernatant after incubation with excess peptide, while those labeled “e” are the remaining complex eluted with imidazole. (**D**) NanoDSF analysis of the ternary complex (5 μM) in isolation and in the presence of excess peptide (10 μM).

### Structure of CHD4-C1bC2ab - GATAD2A-CR2 - CDK2AP ^*ΔN*^ ternary complex

Based on these results, we co-expressed and purified a ternary complex comprised of CHD4-C1bC2ab, GATAD2A-CR2, and CDK2AP1^ΔN^ (**Figure 2B**). The CHD4-C1bC2ab construct included CHD4 domains C1b (residues 1395-1519) and C2ab (residues 1693-1902) connected by a four-amino acid linker (GGGS). This design eliminates the long, unstructured linker between domains and adds 92 amino acids to the C-terminal domain compared with the structure previously crystallized by Zhong et al. ^23^ The GATAD2A-CR2 construct (residues 335-486) includes long N- and C-terminal helices in addition to the central helix and Zn-finger domain involved in binding to CHD4-C1bC2a. The CDK2AP1^ΔN^ construct (residues 29-115) represents the shorter N-terminally truncated isoform as described by Modzelewski *et al*.^26^

We purified the complex using split tandem affinity purification tags, with an N-terminal thioredoxin and hexa-histidine tag on CDK2AP1^ΔN^ and a C-terminal Twin-Strep tag on GATAD2A-CR2. Removing the tags with TEV protease yielded a stable ternary complex as verified by mass photometry. The purified complex readily crystallized forming single crystals that diffracted to at least 2.0 Å resolution. Initial phases were determined by molecular replacement using a portion of the AF3 model of the complex. The final model was refined to a resolution of 2.0 Å with R_free_ = 24.5%.

The overall topology of the complex reveals an interesting mode of interaction between GATATD2A-CR2 and CHD4-C1bC2ab. The N- and C-terminal helices of GATAD2A associate to form a large intervening loop containing the Zn finger domain and the short central a-helix. Hence, GATAD2A-CR2 essentially forms a lasso that wraps around and through the junction between the C1b and C2ab domains of CHD4 (**Figure 2B**). The total buried CHD4-C1bC2ab surface area is 4815 Å^2^, comprising separate interactions with CR2 (2861 Å^2^) and CDK2AP1^ΔN^ (1954 Å^2^).

The interaction between CR2 and CHD4-C1bC2ab involves several distinct features. First, the helical mid-region of CR2 (375-414) positions along the surface of CHD4-C1b and is covered by a “lid” formed by the C-terminal helix of CHD4C2ab (residues 1875-1897). CHD4 completely surrounds the CR2helix (**Figures 2B**) with a network of hydrophobic interactions involving branched sidechain residues as well as a bidentate hydrogen bond between CR2(E391)-C2b(R1801) sidechains. Next, the CR2 zinc finger domain packs against a flat surface of CHD4, forming multiple H-bonds and hydrophobic packing interactions (**Figures 2B**). Finally, the loop of the lasso is closed by a coiled-coil interaction between the C- and N-terminal helices of CR2, which then packs against one side of CHD4-C2ab through an extended hydrophobic surface. Notably, a proline-rich segment (PPPKPPAP), immediately following the CR2 N-terminal a-helix, forms a polyproline II helix that contacts CHD4-C2b (**Figures 2B**).

The C-terminal region of CDK2AP1^ΔN^ forms a helix hairpin that packs against CHD4C2b through hydrophobic interactions and a network of hydrogen bonds. This interaction involves the same surface of CDK2AP1^ΔN^ that mediates self-association to form a homodimer.^27^ Therefore, the CDK2AP1 homodimer must dissociate before binding to CHD4-C2b. In addition, a small segment in the N-terminal region of CDK2AP1^ΔN^ domain, previously shown to be an intrinsically disordered region (IDR) in isolation,^27^ forms a short helix that binds to the alternative binding surface of CHD4-C2a (**Figures 2B**). This interface includes proline-pi interactions and hydrogen bonds.

### Peptide binding destabilizes complex formation

To determine whether the macrocyclic peptides destabilize the interaction between the full GATAD2A-CR2 region and CHD4-C1bC2ab, we purified the binary complex between GATAD2A-CR2 and CHD4-C1bC2ab, with a hexahistidine tag on the CHD4 domains. We bound the complex to nickel immobilized metal affinity chromatography (Ni-IMAC) magnetic beads and incubated the beads with excess peptide. Both GATAD2A-CR2-helix and c38.1B.tr peptides eluted GATAD2A-CR2 from the beads, whereas c2.1A did not (**Figure 2C**). This result shows that selectively targeting the CR2-helix-binding pocket on CHD4-C1bC2ab can disrupt the formation of the full complex.

We then tested whether the peptides destabilized the ternary complex by differential scanning fluorometry. As shown in **Figure 2D**, the ternary complex shows a single cooperative transition with a melting temperature of 56.7 °C. The addition of the truncated peptide (c38.1B.tr) broadens the transition, with an apparent melting temperature of 55.3 °C (ΔT_m_ = -1.4 °C). In contrast, peptide c2.1a, which is predicted to bind to the CDK2AP1 binding surface on CHD4-C2b (Site B), does not modify the observed melting temperature.

## Discussion

In these studies, we sought to identify macrocyclic peptide inhibitors of the interaction between GATAD2A-CR2 and the C-terminal region of CHD4, with the goal of developing a much needed and novel therapeutic strategy to inhibit gene silencing by NuRD. However, the C-terminal region of CHD4 has been difficult to express and purify in isolation for structural and biophysical analyses. A breakthrough came from work by the Mackay group, who developed an expression construct for the C1b and C2a domains of CHD4 by eliminating much of the long (∼150 amino acids) linker between the two domains.^23^ They showed that this region forms a SANT-SLIDE domain, binds DNA, and auto-inhibits CHD4-mediated chromatin remodeling. Studies have also shown that the GATAD2A-CR2 domain binds to this same C-terminal region of CHD4.^16,22^

Building on this prior work, we established an mRNA screen against the CHD4-C2bC1a domains with a peptide library containing fixed cysteine residues cyclized with DBX. The selection produced high-affinity inhibitors characterized by a common C-terminal motif with an additional internal cysteine residue (WCXLLC). Analyzing the enriched sequences using t-distributed stochastic neighbor embedding (t-SNE) reveals that peptides containing this motif form a distinct grouping of clusters (**Figure 1C**) that together represent the majority of the observed clones. Notably, comparing peptide sequences within and between clusters containing this motif does not reveal any clear preferences for the N-terminal portion of the peptides. Accordingly, a truncated peptide comprised of this isolated motif (c38.1B.tr) bound with similar high affinity. Therefore, this small macrocyclic helical peptide likely binds to the same wedge-shaped groove occupied by CR2-helix, competitively disrupts binding by CR2, and destabilizes the ternary complex.

Additionally, the results of the screen highlighted a second potential binding surface on CHD4-C1aC2b. This surface is relatively flat and hydrophobic, lined by multiple phenylalanine (F1765, F1774, and F1781) and leucine (L1711 and L1788) residues. This observation led us to investigate whether CDK2AP1 binds to this alternative site. Co-expressing CDK2AP1 with CHD4-C1bC2ab and GATAD2A-CR2 yielded a stable complex that readily purified. Notably, we had been unable to purify the full GATAD2A-CR2 domain or the CHD4-C1bC2ab domains in isolation despite multiple attempts. Therefore, CDK2AP1 appears to have stabilized the complex and buried hydrophobic surfaces that cause aggregation and proteolytic degradation of the isolated domains.

The crystal structure revealed that CDK2AP1 binds to a long helix hairpin at the C-terminus of CHD4 to form a four-helix bundle, while GATAD2A-CR2 binds the opposite face of the same CHD4 helix hairpin. After this helix hairpin and a short loop, the C-terminus of CHD4 then forms the small helix that binds to the GATAD2A-CR2 helix, forming the lid described above. This C-terminal region of CHD4 is critical for NuRD-dependent silencing of fetal hemoglobin. In a high-throughput mutagenesis study of NuRD in fetal hemoglobin regulation, Sher *et al*.^22^ showed that mutations in this C-terminal region (amino acids 1872-1881), which immediately follows the hairpin helix, do not negatively impact cell fitness while strongly inducing fetal hemoglobin. Hence, CDK2AP1 appears to stabilize a region that defines distinct roles of CHD4 in gene regulation. Consistent with this observation, Bode *et al*.^28^ characterized two distinct NuRD complexes that differ by whether they contain CDK2AP1 and SALL4. Importantly, less CHD4 co-purified in the NuRD complex that lacked CDK2AP1, consistent with CDK2AP1’s role in stabilizing its interaction with GATAD2A-CR2.

CDK2AP1 was first identified as a tumor suppressor frequently lost in oral cancer ^29^ and later shown to associate with cyclin-dependent kinase 2 (CDK2).^30^ More recent work has shown that it plays a key role in the competition between NuRD and the Switch/Sucrose Non-Fermentable (SWI/SNF) chromatin remodeling complex.^31,32^ Incorporation of CDK2AP1 drives a specific function of the NuRD in the mesenchymal to epithelial transition. Interestingly, the structure of the ternary complex shows that CDK2AP1 binds as a monomer via the same interface used for homo-dimerization. The Montelione group^27^ previously determined the structure of CDK2AP1, demonstrating that it forms a homo-dimer by analytical ultracentrifugation, with no evidence of monomeric species at concentrations as low as 10 μM. This observation suggests that dimerization may control whether CDK2AP1 binds to NuRD, thereby shifting its role between regulating the cell cycle by targeting CDK2 for degradation and regulating gene expression by stabilizing CHD4 in NuRD.

As a result, the C-terminal region of CHD4 acts as an interaction hub that dictates NuRD function. The CHD4-C1bC2ab domain binds DNA,^23^ GATAD2A-CR2, and CDK2AP1 through distinct interfaces to control nucleosome remodeling and recruitment of CHD4 to the complex. In these studies, we have identified macrocyclic peptides that target two distinct binding surfaces of CHD4-C1bC2ab. These peptides have the potential for selectively disrupting gene silencing by NuRD without reducing cell fitness. Therefore, these studies shed new light on the structure and function of NuRD, demonstrate that the CHD4-GATAD2A interaction is targetable, and uncover a potential strategy to inhibit NuRD function for therapeutic benefit.

## METHODS

### Materials

All oligonucleotides were purchased from Integrated DNA Technologies (Coraville, IA) or Twist Bioscience (San Francisco, CA). Enzymes Q5 DNA polymerase, PNK, DnpI, and T4-DNA ligase were purchased from New England Biolabs (Ipswich, MA). PURExpress (-aa, -tRNA, -RF123) (E6850Z) translation kits were purchased from New England Biolabs and used as directed for translation of mRNA displayed peptides. Protected amino acids and H-Rink amide resin for solid phase peptide synthesis (SPPS) were purchased from Chem-Impex International. Alexa Fluor^™^ 647 C_2_ Maleimide (A20347) was purchased through Thermo Fisher. EZ-Link^™^ Maleimide-PEG11-Biotin (21911) and chloroacetic anhydride (AC404420250) were purchased from Thermo Fisher Scientific. TR-FRET reagents LANCE Eu-W1024 Streptavidin (AD0063), LANCE Eu-W1024 Anti-6xHis (AD0402) and LANCE Ultra ULight^™^ – anti-GST (TRF0104-D) were purchased from PerkinElmer. HA Tag Antibody conjugated to HRP (A190-108P) was purchased from Bethyl Laboratories, Inc. and SuperSignal^™^ ELISA Pico Chemiluminescent Substrate (37070) was purchased from Thermo Fisher Scientific.

### mRNA display and selection protocols

mRNA display was conducted as previously described. Briefly here, DNA libraries of 10-12 NNK randomized codons were transcribed into RNA and linked to puromycin using Y-ligation with T4 RNA ligase I (NEB, M0204S). The libraries were translated for 30 minutes at 37 °C using a custom GeneFrontier Corporation PUREfrex 2.0 kit (PFC-Z136-2-EX, -aa and -RF123) per the manufacturer specifications. RNA-peptide fusions were reverse transcribed with mMLV reverse transcriptase H (-) (Promega, M3681, 40x) at 42 °C for 1 hour. After reverse transcription, libraries were treated with 100 mM HEPES buffer (pH 7.5), 300 µM NaCl, 0.5 mM TCEP, and 3 mM DBX (α,α′-dibromo-m-xylene; stock solution prepared in ACN), with the final reaction mixture supplemented to 8% ACN. The reaction was incubated at room temperature for

30 min. The displayed library was diluted 10-fold and then incubated with anti-HA magnetic beads (ThermoFisher catalog number: 88836) at a 4:1 ratio of bead slurry to the initial IVT volume. The bead slurry was rotated at room temperature for 30 minutes, followed by three washes with TBS-T (20 mM Tris-HCl, pH 7.5, 150 mM NaCl, 0.1% Tween-20) three times. Next, 4 mg/mL HA peptide in 2x selection buffer (50 mM HEPES, pH 7.5, 300 mM NaCl, 1 mM fresh TCEP, 0.1% Tween-20, 2 mg/mL BSA, 2 mg/mL yeast RNA) was added to the beads, and fusions were eluted by rotating at room temperature for 1 hour. The HA elution volume should be calculated to ensure that the immobilized protein is at a concentration of 200 nM during the selection (50 μL purified library and 250 μL for R1 selection). The resulting libraries were counter selected against magnetic Dynabeads^™^ M-280 Streptavidin (Thermo Fisher Scientific, 110205D) prior to affinity selection against 200 nM immobilized CHD4-C1bC2a. Selections were conducted at 4 °C for 30 minutes followed by washing. Recovered cDNA was eluted by heating at 95 °C for 5 minutes prior to supernatant collection. Collected cDNA was amplified by PCR for preparation of subsequent rounds of selection and NGS analysis. Sequences were translated by their open reading frame and analyzed for enriched families using in-house python scripts.

### General solid phase peptide synthesis procedures (Fmoc-SPPS)

*Fmoc-SPPS*. Peptides were synthesized on 50 mg of H-Rink amide ChemMatrix® resin (loading capacity = 0.48 mmol/g). The resin was swollen for 1 hour in DMF (1.5-3 mL) at room temperature. Coupling reactions were carried out with standard fluorenylmethyloxycarbonyl (Fmoc) synthetic procedures in 10 mL reaction vessels. Coupling reactions were conducted with 5 equivalents of Fmoc-AA-OH, 5 equivalents of HCTU, and 10 equivalents of DIPEA dissolved in 740 µL DMF. Reagents were mixed at room temperature for 15-30 minutes before the reaction vial was drained and washed 4x with DMF and 4x with DCM. Coupled residues were deprotected with the addition of 1 mL 20 % piperidine in DMF. Deprotection reactions were mixed at room temperature 10-15 minutes before the reaction vessel was washed 4x with DMF and 4x with DCM.

#### Acetic anhydride coupling

Acetic anhydride was coupled by adding to the reaction vessel 1 mL of 0.88 M acetic anhydride (36 equiv.) and 0.6 M triethylamine (25 equiv.) in DCM:DMF (1:1). The reaction was mixed at room temperature for 1 hour before washing and drying the resin.

#### Chloroacetic anhydride coupling

Chloroacetic anhydride was coupled by dissolving 100 mg of chloroacetic anhydride (24 equiv.) in 590 µL DCM:DMF (1:1). The solution was added to the reaction vessel and mixed. While mixing, 150 µL (44 equiv.) of TEA was added. The reaction was mixed at room temperature for 2-5 minutes before washing and drying the resin.

#### P*eptide cleavage from resin*

Dried resin was cleaved in standard cleavage cocktail (TFA:TIPS:H_2_O, 95:2.5:2.5) while mixing for 1 hour at 37 °C. The resin was filtered out and TFA evaporated over nitrogen. Resulting peptides were precipitated with 13-15 mL chilled diethyl ether and centrifuged at 4,000 rpm for 10 minutes. The supernatant was decanted, and the peptide pellets dried briefly. Linear peptides were dissolved in minimal DMSO and diluted in ACN:H_2_O (1:1) before purification by HPLC. Thioether bridged peptides were cyclized at this step prior to HPLC purification.

#### DBX (α,α′-Dibromo-m-xylene) Cyclization

After cleavage, the crude linear peptide was dissolved in 500 μL of DMSO and added to a pre-cyclization reaction mixture consisting of 1× PBS buffer (without Tween-20) and acetonitrile (1:1, v/v) to a final volume of 5 mL, containing 10 mM K_2_CO_3_ and 1 mM TCEP. The reaction mixture was rotated at room temperature for 15 min. Cyclization was initiated by the addition of α,α′-dibromo-m-xylene (3.4 mM final concentration; 4.5 mg dissolved in 20 μL of acetonitrile), and the pH was adjusted to >7 using 1 M K_2_CO_3_. The reaction mixture was rotated at room temperature for an additional 30 min. Upon completion, both the pre-cyclization and cyclized peptide samples were analyzed by MALDI-MS, and the cyclized peptides were subsequently purified by reverse-phase preparative HPLC.

### Protein Expression and Purification

*GATAD2A-CR2helix-TAMRA*. DNA encoding for the internal helix (residues 371-404) from the GATAD2A-CR2 domain was cloned into the pET32a(+) vector with N-terminal thioredoxin and hexahistidine tags and a C-terminal TwinStrep tag. A tobacco etch virus protease site (ENLYFQG) immediately precedes the helix such that cleavage with TEV protease removes the N-terminal tags, leaving an additional GS at the N-terminus. A single cysteine residue was included immediately after the CR2-helix. For protein expression, the resulting vector was transformed into Rosetta2(DE3), grown under ampicillin selection at 37 °C until an optical density at 600 nm of approximately 0.9, and induced with 1 mM isopropyl β-d-1-thiogalactopyranoside (IPTG) for ∼3 hours.

The bacteria from 1 L of culture were lysed by sonication in 30 mL of lysis buffer (20 mM Tris pH 8.0, 1 M NaCl, 1 mM dithiothreitol (DTT)) supplemented with 1 mM phenylmethylsulfonyl fluoride (PMSF). The protein was purified by nickel affinity chromatography, eluted with elution buffer (20 mM Tris pH 8.0, 0.2 M NaCl, 1 mM DTT, 300 mM imidazole), digested with TEV protease overnight, and purified over streptactin resin (5 mL Streptactin XT 4Flow, IBA Lifesciences), eluting with biotin elution buffer (BXT, IBA Lifesciences). The protein was concentrated and purified by size-exclusion chromatography (Superdex 75 26/60, Cytiva) equilibrated in TBS (20 mM Tris pH 8.0, 150 mM NaCl) supplemented with 1 mM TCEP. The eluted protein was concentrated and quantified by UV spectroscopy (*E*_(280)_ = 12490 M^-1^ cm^-1^).

The purified peptide was labeled with TAMRA maleimide, 5-isomer (Lumiprobe) following the procedure described by Kim *et al*.^1^. In brief, the peptide was precipitated with ammonium sulfate (70%), resuspended in TBS with a 5-fold excess of TAMRA maleimide, and incubated overnight at room temperature. The labeled peptide was purified by size-exclusion chromatography (Superdex 75 26/60, Cytiva) equilibrated in 25 mM BisTris pH 7.2, 100 mM NaCl, 1 mM DTT, and 50 μM ZnSO_4_ and concentrated. The labeling efficiency and total concentration were quantified by UV spectroscopy (*E*_(558)_ = 84,000 M^-1^ cm^-1^, *E*_(280)_ = 12,490 M^-1^ cm^-1^), and the protein was stored at -80 °C until further use

*CHD4-C1bC2a*. We subcloned the C1b (residues 1395-1519) and C2a (residues 1693-1810) domains of human CHD4 protein, connected with a four amino acid (GGGS) linker, into a modified pET32a (+) with N-terminal thioredoxin and hexahistidine tags and TEV protease site, as described for the GATAD2A-CR2helix. For protein expression, the resulting vector was transformed into Rosetta2(DE3), grown under ampicillin selection at 37 °C until an optical density at 600 nm of approximately 0.6, then the temperature was lowered to 20 °C, and the culture was induced with 0.4 mM isopropyl β-d-1-thiogalactopyranoside (IPTG) overnight (16-20 hours). The bacteria from 1 L of culture were lysed by sonication in 30 mL of buffer A (20 mM Tris pH 8.0, 0.5 M NaCl, 1 mM dithiothreitol (DTT), 5% glycerol) supplemented with 1 mM phenylmethylsulfonyl fluoride (PMSF). The protein was purified by nickel affinity chromatography, eluted with buffer A + 300 mM imidazole, digested with TEV protease while dialyzing against buffer A overnight, and re-purified by nickel affinity chromatography, collecting the flow-through. The protein was concentrated and purified by size exclusion chromatography (Superdex 75 10/300 Increase, Cytiva) in buffer A. The eluted protein was concentrated, quantified by UV spectroscopy (*E*_(280)_ = 43430 M^-1^ cm^-1^), and stored at -80 °C until further use.

For biotinylated samples, the expression construct was modified to include a C-terminal avi-tag (GGLNDIFEAQKIEWHEG) and the protein purified as above. We bacterially expressed and purified BirA enzyme to biotinylate CHD4-C1bC2a-avi *in vitro*.^2^ pET21a-BirA was a gift from Alice Ting (Addgene plasmid # 20857 ; http://n2t.net/addgene:20857; RRID:Addgene_20857). Biotinylation was confirmed by a streptavidin gel shift assay.^3^

#### Binary Complex (CHD4-C1bC2ab + GATAD2A-CR2)

We subcloned the CR2 domain (amino acids 335-486) of the human GATAD2A protein into the pET28a vector with an N-terminal maltose-binding protein (MBP) and a C-terminal TwinStrep tag. A TEV protease site was included before and after the CR2 domain such that cleavage with TEV protease leaves two additional residues (GS) on the N-terminus and an additional 13 residues on the C-terminus (GSGGSSAENLYFQ). We subcloned the C1b (residues 1395-1519) and C2ab (residues 1693-1902) domains of human CHD4 protein, connected with a four amino acid (GGGS) linker, into the pCDF vector with N-terminal thioredoxin and hexahistidine tags and a TEV protease site as described for CHD4-C1bC2a.

For protein expression, the resulting vectors were co-transformed into Rosetta2(DE3), grown under kanamycin and streptomycin selection at 37 °C until an optical density at 600 nm of approximately 0.6, the temperature was lowered to 20 °C, and the culture induced with 0.4 mM isopropyl β-d-1-thiogalactopyranoside (IPTG) overnight (16-20 hours). The bacteria from 1 L of culture were lysed by sonication in 30 mL of buffer A supplemented with 1 mM PMSF. The complex was purified by sequential affinity chromatography over nickel (5 mL HisTRAP, Cytiva), streptactin (5 mL Streptactin XT 4Flow, IBA Lifesciences), and amylose (5 mL MBPTrap, Cytiva) resins, eluting with 300 mM imidazole, 5 mM biotin, and 10 mM maltose, respectively. The complex was concentrated and purified by size exclusion chromatography (Superdex 75 Increase 10/300, Cytiva) in buffer A. The eluted complex was concentrated, quantified by UV spectroscopy (*E*_(280)_ = 147710 M^-1^ cm^-1^), and stored at - 80 °C until further use.

#### Ternary Complex (CHD4-C1bC2ab + GATAD2A-CR2 + CDK2AP1)

We subcloned the same CR2 domain from GATAD2A into the first multiple cloning site of the pETDuet-1 vector, with N-terminal Flag and TwinStrep tags and an intervening TEV protease site. We subcloned the same C1bC2ab domains of CHD4 with a GGGS linker into the second multiple cloning site of the same pETDuet-1 vector, without any affinity purification tags. We subcloned the short isoform of CDK2AP1 (amino acids 29-115) into the pCDF vector, with an N-terminal thioredoxin and hexahistidine tag and a TEV protease site, as described above.

For protein expression, the resulting vectors were co-transformed into Rosetta2(DE3), grown under ampicillin and streptomycin selection at 37 °C until an optical density at 600 nm of approximately 0.6, then the temperature was lowered to 20 °C, and the culture was induced with 0.4 mM IPTG overnight (16-20 hours). The bacteria from 1 L of culture were lysed by sonication in 30 mL of buffer A supplemented with 1 mM PMSF. The complex was purified by sequential affinity chromatography over nickel (5 mL HisTRAP, Cytiva) and streptactin (5 mL Streptactin XT 4Flow, IBA Lifesciences) resins, eluting with 300 mM imidazole and 5 mM biotin, respectively. The affinity tags were removed by TEV protease cleavage overnight at room temperature. The complex was then re-purified over the nickel column, collecting the flow-through, and concentrated and purified by size-exclusion chromatography (Superdex 75 Increase 10/300, Cytiva) in buffer A. The eluted complex was concentrated, quantified by UV spectroscopy (*E*_(280)_ = 60850 M^-1^ cm^-1^), and stored at -80 °C until further use.

### Fluorescence Polarization Assays

*Binding assay*. CHD4-C1bC2a was serially diluted in buffer A with 40 nM of TAMRA-GATAD2A-CR2helix. Fluorescence polarization was measured on a CLARIOstar microplate reader (BMG Labtech), and the observed polarization fit to a general equation for two-state binding to determine the dissociation constant (K_D_) and plotted using Prism software (GraphPad).

#### Competitive Inhibition Assay

Unlabeled peptides were serially diluted in buffer with 40 nM of TAMRA-GATAD2A-CR2helix and 100 nM of CHD4-C1bC2a. The observed polarization was fit to an explicit equation for competitive inhibition^4^ as implemented by Lee *et al*.^5^ to determine the inhibitor dissociation constant (K_D_^inhib^). For those peptides that showed non-competitive binding (c2.1A), the observed polarization was fitted to the two-state binding equation described above. For comparison between datasets, the resulting polarization was normalized to range from 0 to 100.

### AlphaFold 3 models

We generated models of the CHD4-C1bC2a domain bound to macrocyclic peptides with AphaFold 3.0 implemented on a local computer cluster with an L40 48 GB GPU (NVIDIA). Peptide cyclization was modeled using as a linker 1,3 dimethylbenzene (CCD Code: 8VH) with bonds between the methyl carbons in the linker and two cysteine sulfur atoms in the respective peptides. Models of CHD4-C1bC2ab bound to GATAD2A-CR2 were generated on the AlphaFold 3 server (alphafoldserver.com).

### NanoBRET Assays

The C1bC2ab domains (1395-1551, 1690-1902) and C1bC2a (residues 1395-1551, 1690-1810) were cloned into the pNLF1-N and pNLF1-C vectors (Promega) in frame with an N-or C-terminal NanoLuciferase fusion, respectively. CDK2AP1 full-length (1-115) and isolated helix-turn-helix (60-115) were cloned into the pHTC vector (Promega) in frame with a C-terminal HaloTag domain. Intracellular NanoBRET was measured following manufacturers protocol as described previously.^6^

### Surface-based biophysical measurement of binding kinetics

Grated-Coupled Interferometry (GCI) experiments were performed on a Creoptix WAVE system (Creoptix, AG) using 4PCH STA WAVE sensor chips (polycarboxylate surface, streptavidin coated). Chips were conditioned with borate buffer (100 mM sodium borate pH 9, 1 M NaCl). Biotinylated CHD4-C1bC2a was directly immobilized onto the sensor chip by injection onto the surface in Tris [20 mM Tris-HCl pH 8.0, 500 mM NaCl, 1 mM TCEP (fresh, pH 7.0), 0.05% Tween-20] buffer at 10 μg/mL at a flowrate of 10 μL/min. Biotinylated protein was immobilized onto a streptavidin WAVEchip® (Type: 4PCH-STA) to a surface density of ∼4000 response units (pg/mm^2^). Kinetic analyses were performed at 25°C using traditional multicycle kinetics where the analyte is introduced at increasing concentrations with injection of uniform duration. The analyte was injected over a 7–12-point, three-fold serial dilution series with varying maximum concentrations (starting at 5 µM peptide; concentrations exhibiting poor fitting curves were excluded). Injections were performed at a flow rate of 15 µL/min per channel, with a 60 s association phase followed by a 60 s dissociation phase. Binding sonograms were generated using a standard 1:1 binding kinetics model and kinetics constants determined using standard WAVEcontrol evaluation software.

### Circular dichroism (CD) analysis

Lyophilized peptides (powder) were initially dissolved in 1:1 ACN/H2O. The concentration was determined by UV/Vis spectroscopy based on the molecular weight and molar absorptivity for each peptide. Then peptides were diluted to 100 µM in 10 mM sodium phosphate buffer pH 7.5 and 1% ACN. CD spectra were acquired using an Jasco J816 CD spectrometer. Measurements were acquired with taken every 0.5 nm at scanning speed of 50 nm/min from wavelengths of 250 nm to 190 nm. A high precision cell with 0.1 mm light path (Hellma Analytics, Art. No. 1101-0.1-40) was used to house the samples. Three spectra acquisitions were taken for each sample. A blank comprised of 1% ACN in 10 mM sodium phosphate buffer pH 7.5 was first run followed by each peptide (c38.1B and c16.1B). The blank values were subtracted from each sample.

### Ternary complex crystallization and structure determination

We set up six crystallization trials of the ternary complex using a Mosquito robot (SPT Labtech) in 96-well Swissci Intelliplates at 25 °C in a sitting-drop format. Each drop contained 0.2 μL of complex (11.8mg/mL, in 20 mM Tris pH 8, 150 mM NaCl, 1 mM DTT, 0.02% NaN_3_) and 0.1 μL of screening solution. The final crystallization hit was obtained using the MCSG2 screen (Microlytic) in 0.2 M sodium malonate, pH 7.0, and 20% (w/v) PEG 3350. Single crystals were harvested into a cryoprotectant solution with 25% ethylene glycol and cooled in liquid nitrogen. X-ray diffraction data sets were recorded on the APS sector 22 (ID-D) at the University of Georgia at 100 K and a wavelength of 1.0 Å. The data was reduced locally using HKL2000.^7^ We used the Phenix software package for data analysis and refinement,^8,9^ and Phaser ^10^ to determine the initial phases by molecular replacement using an AF 3 model of the complex.

## ASSOCIATED CONTENT

## Supporting Information

The following files are available free of charge:

Supplementary Methods and Figures (PDF)

## AUTHOR INFORMATION

## Author contributions

DCW, AAB, and GDG conceived of the experiments. DCW, JR, and AAB wrote the manuscript. DCW, JR, TL, JMP, and DD performed experiments. DCW, AAB, and RKM analyzed results.

